# Hypoxia Bi-directionally Regulates Gut Vascular Barrier through HIF-1α-dependent Mechanism *in vitro*

**DOI:** 10.1101/2023.05.26.542539

**Authors:** Ping Liu, Wen Dai, Jing Du, Yingshan Zhou, Xiaodu Wang, Houhui Song

## Abstract

**Objective:** The gut vascular barrier (GVB) is the key checkpoint for pathogens to enter the blood circulation through the intestine, which is crucial for maintaining the intestinal barrier function. However, the effect and molecular mechanism of hypoxia on GVB remains unclear. Here, we show a role of the transcription factor hypoxia inducible factor-1α (HIF-1α) in hypoxia-induced bi-directional regulation of GVB.

**Approach and Results:** An *in vitro* GVB model composed of rat intestinal microvascular endothelial cells was studied. Evans blue-albumin efflux assay showed that the experimentally severe hypoxia induced by cobalt chloride (500 μM, 24 h) markedly disrupted the GVB *in vitro*, while mild hypoxia induced by cobalt chloride (500 μM, 6 h) evidently enhanced the GVB, revealing hypoxia-induced bi-directional regulation of the GVB for the first-time. Importantly, knockdown of HIF-1α largely abolished the bi-directional changes of GVB caused by hypoxia. Furthermore, experimentally severe hypoxia exacerbated the inflammatory GVB disruption induced by LPS or TNF-α, while the mild hypoxia promoted the repair.

**Conclusion:** Collectively, our data indicate that hypoxia bi-directionally regulates GVB in a HIF-1α-dependent manner.

## Introduction

Gut vascular barrier (GVB), the last guard between intestine and blood circulation, is a structured endothelium below the intestinal epithelial layer that plays a fundamental role in avoiding indiscriminate trafficking of bacteria which has escaped from the epithelial barrier ^1,2^. GVB works together with intestinal epithelial barrier (IEB) in dynamic maintenance of intestinal barrier homeostasis^3,4^. Imbalance of GVB leads to pathogen translocation, leakage of body fluid and intestinal microenvironment disorders, mediating diseases of the gut-liver axis^1,2,4–6^ and gut-brain axis^3,7,8^, as well as colorectal cancer (CRC) metastasis^9–11^. Thus, understanding key events and regulators governing GVB, especially in the context of intestinal microenvironments, would deepen our understanding of basic biology of GVB and aid in designing therapeutic approaches targeting GVB.

Hypoxia is a significant component of the inflammatory microenvironment within the intestinal mucosa^12–14^. The intestine, as a mucosal organ, is supported by a rich and complex underlying vasculature. The anatomy and function of the intestine provide a fascinating oxygenation profile as, even under physiologic conditions, the intestinal mucosa experiences profound fluctuations in blood flow and metabolism^12^. Also, in inflammatory conditions such as inflammatory bowel diseases (IBD), the increased oxygen demand by resident and gut-infiltrating immune cells coupled with vascular dysfunction brings about a marked reduction in mucosal oxygen concentrations^13,15,16^. For this reason, the intestine, particularly the barrier-protective GVB, are much more susceptible to damage related to diminished blood flow and concomitant tissue hypoxia.

The transcription factor hypoxia inducible factor (HIF) −1 functions as a master regulator of oxygen homeostasis^15,17^. Previous studies *in vitro* and *in vivo* have demonstrated that the activation of epithelial HIF serves as an alarm signal for the resolution of inflammation in various murine disease models^12,16,18,19^. Nevertheless, until now, how GVB adapt and respond to intestinal hypoxic microenvironments remain surprisingly limited. Here, we sought to identify compensatory mechanisms that regulate GVB under hypoxic microenvironment. Our investigation reveals a bi-directional regulatory mechanism whereby the HIF-1α orchestrates the function of GVB, which is different from that of the IEB.

## Results

### Distinguishing sensitivity of the intestinal microvascular endothelial cells and the intestinal epithelial cells under hypoxia

First, we tested the effects of hypoxia on the viability of the rat intestinal microvascular endothelial cells RIMVEC-11 and the intestinal epithelial cells IEC-6, comparatively. The hypoxic cellular models were chemically induced by cobalt chloride (CoCl_2_). As shown in Figure 1A, at 6 h or 12 h time point, none of 100, 200, 300, 400, 500 and 600 μM CoCl_2_ treatments showed any significant effect on the viability of RIMVECs; at 24 h time point, with the increase of CoCl_2_ induced concentration, the cell viability of RIMVECs showed a significant trend of increasing (200, 300 and 400 μM CoCl_2_, *P* < 0.05) and then decreasing (600 μM CoCl_2_, *P* < 0.05), with neither 100 μM nor 500 μM showing significant effects on the viability of RIMVECs; similarly, at 48 h time point, with the increase of CoCl_2_ concentration, the cell viability of RIMVECs showed a significant trend of increasing (200 and 300 μM CoCl_2_, *P* < 0.05) and then decreasing (500 and 600 μM CoCl_2_, *P* < 0.05), with neither 100 μM nor 400 μM showing any significant effect on the viability of RIMVECs. Meanwhile, under light microscope (Figure 1C & EV1A), with the increase of hypoxia induction for 6 h, 12 h, and 24 h, RIMVECs gradually showed an evident trend of morphological loss, volume reduction and enlarged intercellular gaps (especially with 400, 500, and 600 μM CoCl_2_ for 24 h). By comparison, as shown in Figure 1C, at 6 h, 12 h and 24 h time point, none of 100, 200, 300, 400, 500 and 600 μM CoCl_2_ treatments showed any significant effect on the viability of IECs; at 48 h time point, with the increase of CoCl_2_ concentration, the cell viability of IECs showed a significant trend of increasing (100, 200 and 300 μM CoCl_2_, *P* < 0.05) and then decreasing (600 μM CoCl_2_, *P* < 0.05), with neither 400 μM nor 500 μM showing any significant effect on the viability of IECs. Moreover, under light microscope (Figure 1C & EV1A), with the increase of hypoxia induction for 6 h, 12 h and 24 h, IECs cells did not show any evidently morphological changes. These data indicate that the intestinal microvascular endothelial cells are distinguishingly more susceptible to experimental hypoxia than the intestinal epithelial cells are *in vitro*. Meanwhile, 100 μM and 500 μM dosage of CoCl_2_ induction for 6 h or 24 h respectively that were noncytotoxic were used in subsequent experiments.

**Figure 1.**
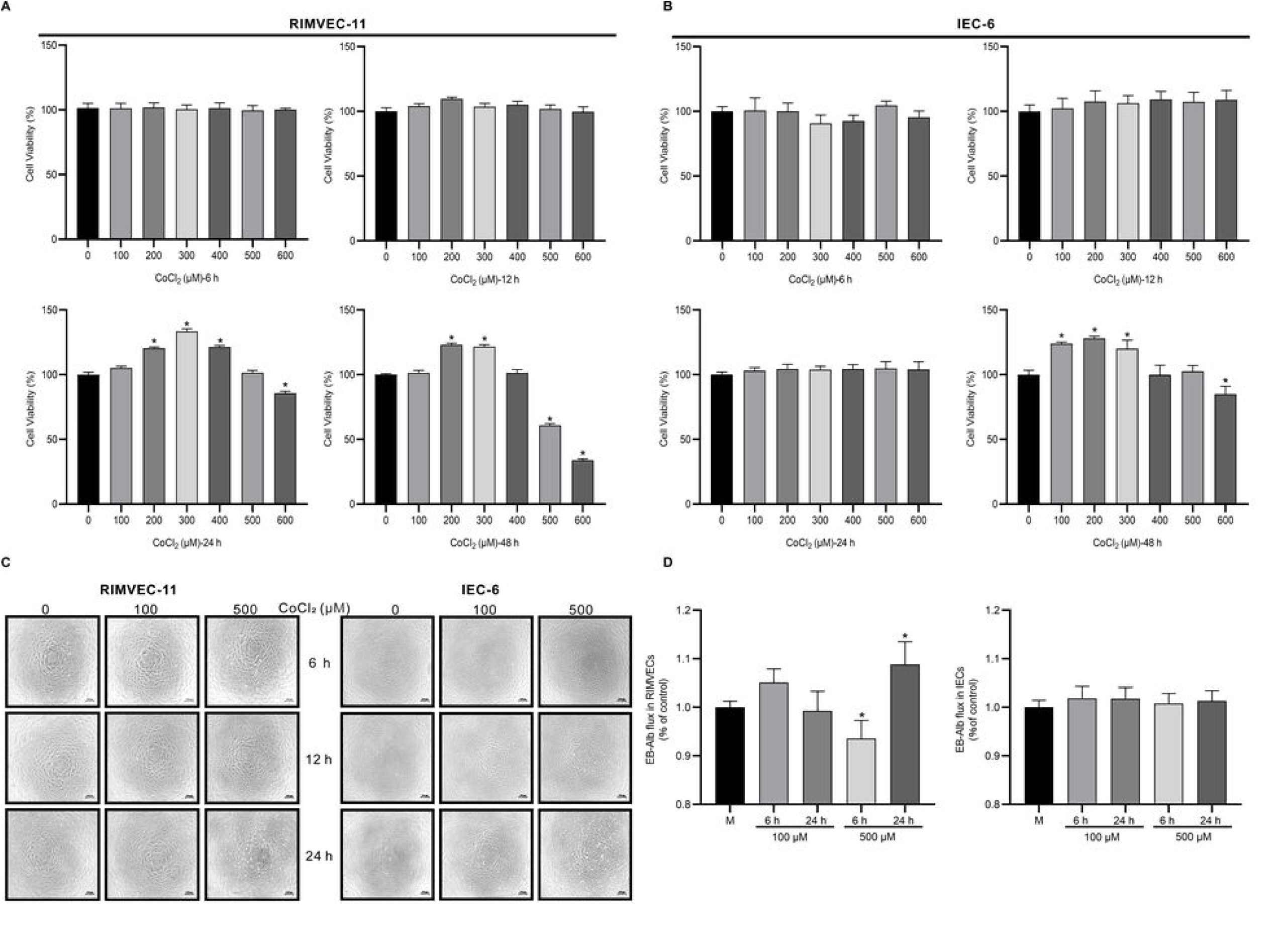
Distinguishing sensitivity of the intestinal microvascular endothelial cells and bi-directional regulation *in vitro* of GVB under hypoxia. (A) The cell viability of RIMVEC-11 treated with CoCl_2_ (0 - 600 μM) for 6 h, 12 h, 24 h, and 48 h by CCK-8 assay (n = 6). (B) The cell viability of IEC-6 treated with CoCl_2_ (0 - 600 μM) for 6 h, 12 h, 24 h, and 48 h by CCK-8 assay (n = 6). (C) The cell morphology of RIMVEC-11 and IEC-6 treated with CoCl_2_ (0 μM, 100 μM, and 500 μM) for 6 h, 12 h, and 24 h. Scale bars = 100 μm. (D) Effects of different degrees of hypoxia (100 μM CoCl_2_ for 6 h or 24 h; 500 μM CoCl_2_ for 6 h or 24 h) on the permeability of *in vitro* GVB (composed of RIMVEC-11) and *in vitro* IEB (composed of IEC-6) (n = 3). The permeability of the cell monolayer barriers was determined by EB-albumin efflux assay. Data information: In (A, B and D), data are presented as mean ± SD. \**P* < 0.05 was compared with the control group.

### GVB responses bi-directionally to hypoxia *in vitro*

To investigate the differential effects of hypoxia on GVB and IEB *in vitro*, we analyzed the transcription factor HIF-1α at transcript level by quantitative PCR both in RIMVECs and IECs (Figure EV1B). In RIMVECs, HIF-1α increased significantly (*P* < 0.05) under hypoxia (100 μM CoCl_2_ for 6 h or 24 h; 500 μM CoCl_2_ for 6 h or 24 h). Specifically, under 100 μM CoCl_2_ induced hypoxia, HIF-1α in RIMVECs showed a significant trend of increasing at 6 h and then sharply decreasing at 24 h; under 500 μM CoCl_2_ induced worse hypoxia, HIF-1α in RIMVECs showed a significant trend of increasing at 6 h, while then even more sharply increasing at 24 h. By comparison, in IECs, the tendency of HIF-1α alteration at mRNA levels was similar as in RIMVECs, whereas evidently the degree of alteration in RIMVECs was much greater than that in IECs.

Meanwhile, to determine how hypoxia influence the function of these two barriers, we tested the permeability of *in vitro* GVB composed of RIMVECs and *in vitro* IEB composed of IECs with hypoxia induction using the Evans Blue-albumin assay, respectively. As shown in Figure 1D, in the GVB, 6 h of hypoxia induction (500 μM CoCl_2_) significantly decreased the content of EB-albumin in the lower chambers compared with the control group (*P* < 0.05), which means the experimentally mild hypoxia (500 μM CoCl_2_ for 6 h) could markedly enhance the function of GVB. However, 24 h of hypoxia induction (500 μM CoCl_2_) significantly increased the content of EB-albumin in the lower chambers compared with the control group (*P* < 0.05), suggesting that the experimentally severe hypoxia (500 μM CoCl_2_ for 24 h) could evidently lead the hyperpermeability and barrier disruption of GVB. Meanwhile, 6 h or 24 h of 100 μM CoCl_2_ induction did not show any significant effect on the permeability of *in vitro* GVB functionally. By comparison, in the IEB, none of the chemical hypoxia induction (100 μM CoCl_2_ for 6 h or 24 h; 500 μM CoCl_2_ for 6 h or 24 h) showed any significant effect on the content of EB-albumin in the lower chambers compared with the control group, suggesting that the IEB shows certain evident resistance under hypoxic conditions. Combing the results of the two barriers, the GVB has a specifically bi-directional regulation under hypoxia condition functionally.

### Hypoxia modulates HIF-1α and intercellular tight junction proteins in RIMVECs

To determine whether hypoxia regulates the GVB through HIF-1α and its mediated interendothelial tight junctions (TJs), we analyzed the protein expression (Figure 2A) and cellular distributions (Figure 2B & 2C) of HIF-1α, as well as the main TJ proteins including ZO-1, Occludin and Claudin-1 (Figure 2D) using Western blotting and immunofluorescence assay. As shown in Figure 2A, the protein level of HIF-1α significantly increased (*P* < 0.05) under different degrees of hypoxia (100 μM CoCl_2_ for 6 h or 24 h; 500 μM CoCl_2_ for 6 h or 24 h) in RIMVECs. Through nucleoplasm extraction (Figure 2B), we found that the levels of HIF-1α protein transferred from cytoplasm to nucleus significantly increased (*P* < 0.05) in RIMVECs under different degrees of hypoxia (100 μM CoCl_2_ for 6 h; 500 μM CoCl_2_ for 6 h or 24 h). Notably, with 100 μM CoCl_2_ induction, the level of HIF-1α protein entering nuclear in RIMVECs showed a significant trend of increasing (at 6 h time point, *P* < 0.05), and then decreasing (at 24 h time point, *P* < 0.05) to normal level; while with 500 μM CoCl_2_ induction, the level of HIF-1α protein entering nuclear increased (at 6 h time point, *P* < 0.05), and then maintained (at 24 h time point), which was confirmed by immunofluorescence in Figure 2C. This indicates that HIF-1α is activated to enter nucleus to play a regulatory role in RIMVECs during certain hypoxia.

**Figure 2.**
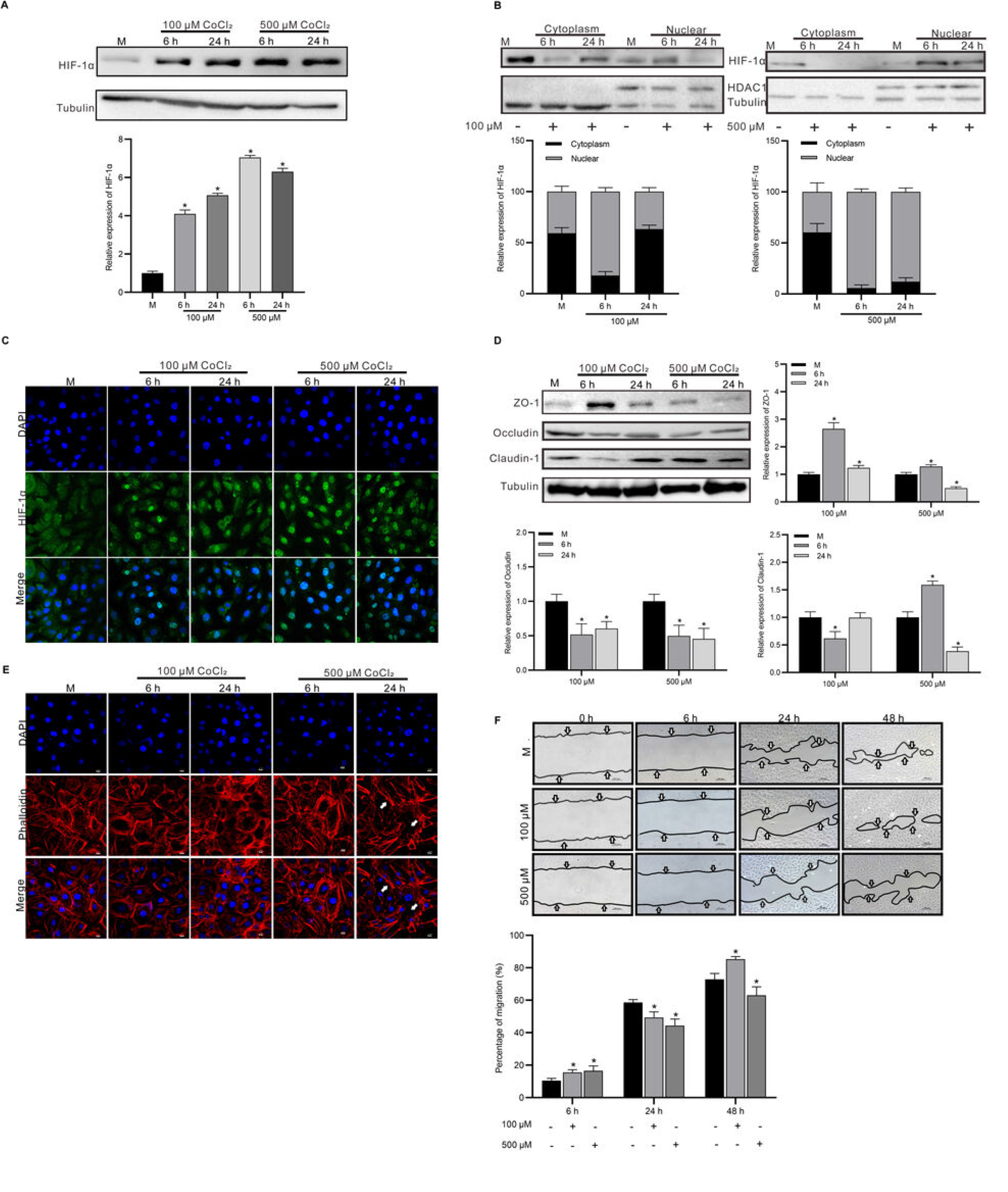
Hypoxia modulates HIF-1α and intercellular tight junction proteins in intestinal microvascular endothelial cells. (A) Western blotting analysis verified the increasing HIF-1α protein in RIMVEC-11 (n = 3). (B) Nuclear protein extraction showed the presence of HIF-1α protein by immunoblotting after 6 h or 24 h exposure to CoCl_2_ (100 μM or 500 μM) in RIMVEC-11. Tubulin and HDAC1 were specific markers for the cytoplasm and nuclear components, respectively (n = 3). (C) Immunofluorescence assay verified the distribution of HIF-1α protein under hypoxia (100 μM CoCl_2_ for 6 h or 24 h; 500 μM CoCl_2_ for 6 h or 24 h) in RIMVEC-11. HIF-1α (green) was co-stained with DAPI (blue). Scale bars = 10 μm. (D) Western blotting analysis verified the expression of tight junction proteins (ZO-1, Occludin, and Claudin-1) under hypoxia (100 μM CoCl_2_ for 6 h or 24 h; 500 μM CoCl_2_ for 6 h or 24 h) in RIMVEC-11 (n = 3). (E) Immunofluorescence assay verified the cytoskeleton changes of RIMVEC-11 under hypoxia (100 μM CoCl_2_ for 6 h or 24 h; 500 μM CoCl_2_ for 6 h or 24 h). Phalloidin (red) was co-stained with DAPI (blue). Scale bars = 10 μm. (F) Scratch assay verified the endothelial migration potential of RIMVEC-11 under hypoxia. Quantification of recovery in scratches under experimental hypoxia (100 μM or 500 μM CoCl_2_) for 0, 6, 24, and 48 h (n = 3). Scale bars = 100 μm. Data information: In (A, B, D and F), data are presented as mean ± SD. \**P* < 0.05 was compared with the control group.

Furthermore, in terms of TJ proteins, as shown in Figure 2D, both 100 and 500 μM CoCl_2_ induction notably promoted first (at 6 h time point, *P* < 0.05) and then inhibited (at 24 h time point, *P* < 0.05) the expression of ZO-1; both 100 and 500 μM CoCl_2_ induction significantly inhibited (at 6 h or 24 h time point, *P* < 0.05) the expression of Occludin; notably, 100 μM CoCl_2_ induction notably inhibited first (at 6 h time point, *P* < 0.05) and then promoted (at 24 h time point, *P* < 0.05) the expression of Claudin-1 to normal level, while 500 μM CoCl_2_ induction markedly promoted first (at 6 h time point, *P* < 0.05) and then inhibited the expression of Claudin-1 (at 24 h time point, *P* < 0.05), which is consistent with the trend of GVB permeability. Together, these results show that the experimental hypoxia could markedly modulate HIF-1α and intercellular tight junction protein in the intestinal microvascular endothelial cells *in vitro*.

### Hypoxia modulates endothelial migration and cell cytoskeleton of RIMVECs

The endothelial migration and cell cytoskeleton of RIMVECs under hypoxia were determined. The intracellular F-actin was assessed using immunofluorescence assay to confirm the alteration of cell cytoskeleton of RIMVECs in GVB (Figure 2E). Meanwhile, we assessed the potentials of endothelial migration of RIMVECs and the repair of GVB injury under hypoxia using scratch assay (Figure 2F). Under hypoxia (100 μM CoCl_2_ for 6 h or 24 h; 500 μM CoCl_2_ for 6 h), compared with the control group, the F-actin was intactly and homogeneously distributed in RIMVECs, without any obviously intercellular gap; while the severe hypoxia (500 μM CoCl_2_ for 24 h) induced the abnormal reorganization and redistribution of cell cytoskeleton, remarkably wrinkled cell morphology and enlarged intercellular gaps. These data suggest that the mild hypoxia maintains and strengthens the GVB structurally at the endothelial level, whereas the severe hypoxia disrupts it. Meanwhile, as shown in Figure 2F, with hypoxia of 100 μM CoCl_2_ induction, the endothelial migration of RIMVECs showed a significant trend of increasing (at 6 h time point, *P* < 0.05), then decreasing (at 24 h time point, *P* < 0.05), and then increasing (at 48 h time point, *P* < 0.05) compared with the control group with increase of time. However, with hypoxia of 500 μM CoCl_2_ induction, the endothelial migration of RIMVECs showed a significant trend of increasing (at 6 h time point, *P* < 0.05) then decreasing (at 24 h time point, *P* < 0.05), and then decreasing (at 48 h time point, *P* < 0.05) with increase of time. These data suggest that the repair potentials of GVB injury could be activated by mild hypoxia, while inhibited by severe hypoxia eventually. Together, these results indicate that hypoxia bi-directionally regulates GVB functionally and structurally *in vitro*.

### Hypoxia regulates GVB through a HIF-1α-dependent mechanism

To elucidate the role of HIF-1α in hypoxia-induced modification of GVB functionally and structurally, we downregulated HIF-1α expression in RIMVEC-11 by shRNA (Figures EV 2), and analyzed the protein expression (Figure 3A) of HIF-1α and Claudin-1 under experimentally mild or severe hypoxia in sh-HIF RIMVEC-11 using Western blotting. As shown in Figure 3A, the HIF-1α knockdown regulated the effects of hypoxia on Claudin-1 and made Claudin-1 protein “blunted” to a certain extent in RIMVECs. Meanwhile, through scratch assay (Figure 3B), the endothelial migration of sh-HIF RIMVECs was determined. Compared with NC group, the scratch healing rate of sh-HIF RIMVECs decreased significantly at 6 h, 24 h and 48 h (*P* < 0.05) with increase of time, suggesting that HIF-1α knockdown markedly results in an evident decrease in endothelial migration of RIMVECs. In addition, under 100 μM CoCl_2_-induced experimental hypoxia, the scratch healing rate of sh-HIF RIMVECs showed a significant trend of decreasing (at 6 h time point, *P* < 0.05), then increasing to equal level (at 24 h time point, *P* > 0.05), and finally decreasing (at 48 h time point, *P* < 0.05) compared with NC group with increase of time; under 500 μM CoCl_2_-induced hypoxia, the endothelial migration of sh-HIF RIMVECs showed a significant trend of decreasing (at 6 h time point, *P* < 0.05), then increasing to equal level (at 24 h time point, *P* > 0.05), and continuously increasing (at 48 h time point, *P* < 0.05) compared with NC group with increase of time. These data suggest that HIF-1α mediates the regulation of hypoxia on GVB injury repair. Furthermore, using the Evans Blue-albumin assay, we tested the permeability of *in vitro* GVB composed of sh-HIF RIMVECs with hypoxia induction. As shown in Figure 4A, compared with NC group, the bi-directional modification of different degrees of experimental hypoxia on the permeability of *in vitro* sh-HIF GVB were all evidently inhibited (*P* < 0.05). These results indicate that hypoxia regulates the GVB functionally through a HIF-1α-dependent mechanism. Using immunofluorescence assay, the endothelial cytoskeleton changes of sh-HIF RIMVECs in GVB were also determined (Figure 4B). Distinctly, compared with NC group, the structurally destructive effects of severe hypoxia (500 μM CoCl_2_ for 24 h) on GVB, including abnormal reorganization and redistribution of cytoskeleton, wrinkled cell morphology and enlarged intercellular gaps, were all eliminated in sh-HIF RIMVECs. These data verify the key role of HIF-1α in hypoxia-induced modification of GVB structurally. Collectively, hypoxia regulates GVB through a HIF-1α-dependent mechanism.

**Figure 3.**
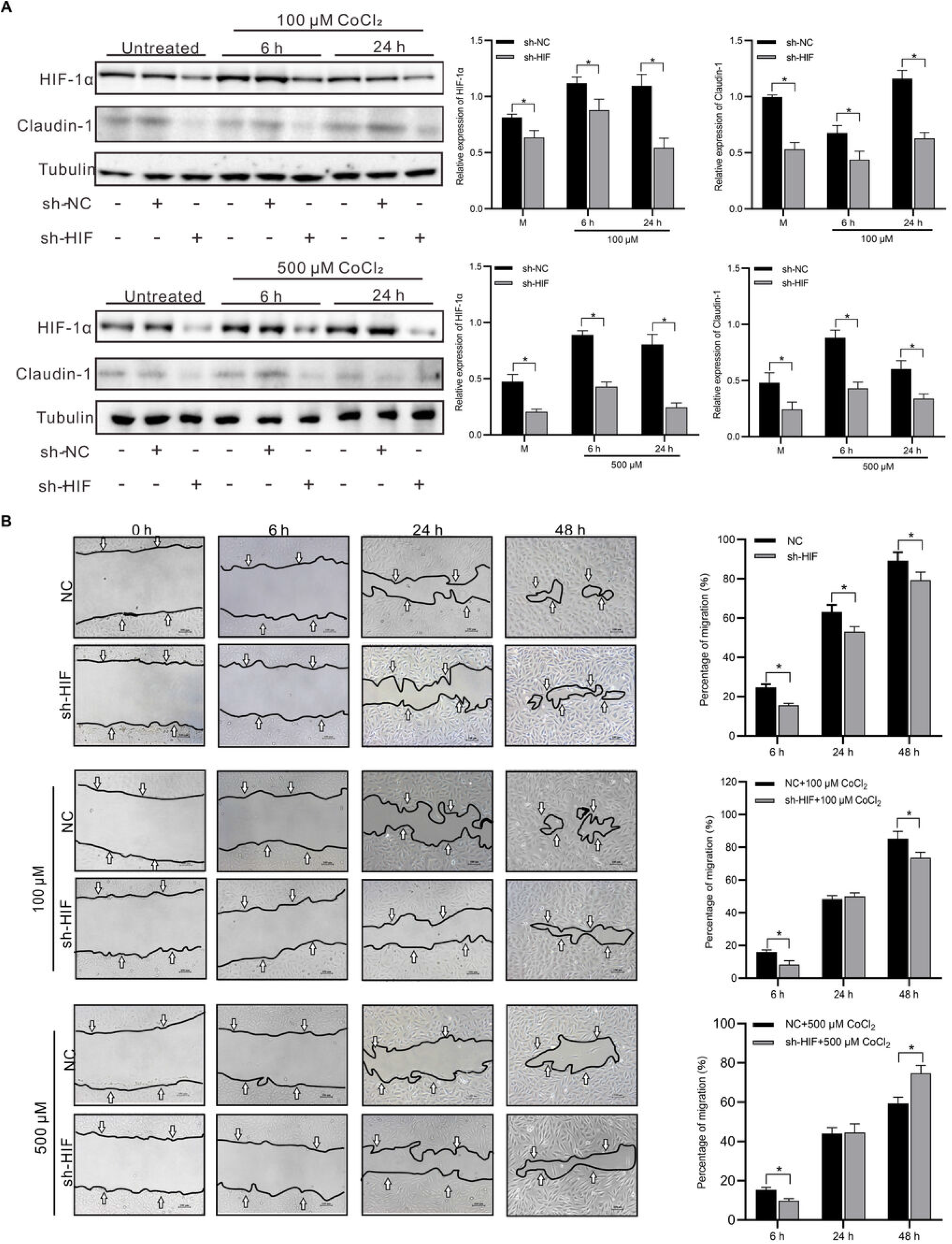
Knockdown of HIF-1α switches hypoxia-induced changes in intestinal microvascular endothelial cells. (A) Western blotting analysis verified the expression of HIF-1α/Claudin-1 under experimentally mild or severe hypoxia (100 μM CoCl_2_ for 6 h or 24 h; 500 μM CoCl_2_ for 6 h or 24 h) in sh-HIF or sh-NC RIMVEC-11 (n = 3) (B) Scratch assay verified the endothelial migration potential of sh-HIF or sh-NC RIMVEC-11 under hypoxia. Quantification of recovery in scratches under experimental hypoxia (100 μM or 500 μM CoCl_2_) for 0, 6, 24, and 48 h (n = 3). Scale bars = 100 μm. Data information: In (A-B), data are presented as mean ± SD. \**P* < 0.05 was compared with the sh-NC group.

**Figure 4.**
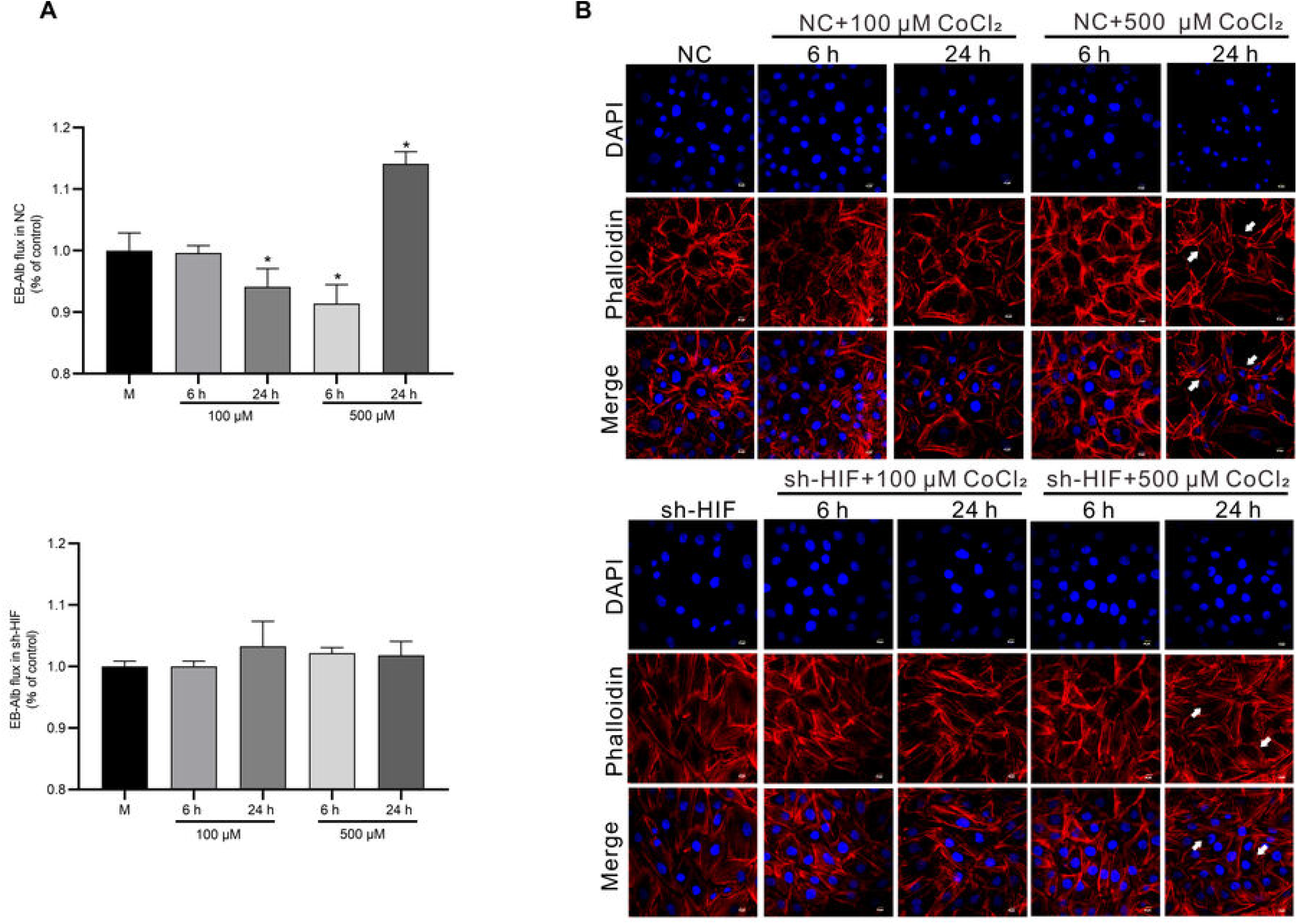
Knockdown of HIF-1α inhibits hypoxia-induced bi-directional alteration of GVB *in vitro*. (A) Effects of experimentally mild or severe hypoxia (100 μM CoCl_2_ for 6 h or 24 h; 500 μM CoCl_2_ for 6 h or 24 h) on the permeability of *in vitro* GVB (composed of sh-HIF or sh-NC RIMVEC-11). The permeability of the cell monolayer barriers was determined by EB-albumin efflux assay (n = 3). (B) Immunofluorescence assay verified the cytoskeleton changes of sh-NC or sh-HIF RIMVEC-11 under experimentally mild or severe hypoxia (100 μM CoCl_2_ for 6 h or 24 h; 500 μM CoCl_2_ for 6 h or 24 h). Phalloidin (red) was co-stained with DAPI (blue). Scale bars = 10 μm. Data information: In (A), data are presented as mean ± SD. \**P* < 0.05 was compared with the sh-NC group.

### The effects of hypoxia treatment on inflammatory GVB disruption

To assess the effects of hypoxia treatment on GVB disruption, the inflammatory GVB disruption models were induced by lipopolysaccharide (LPS, 1 μM) or tumour necrosis factor-α (TNF-α, 0.3 μM) respectively. The protein levels of Claudin-1 in RIMVECs were analyzed by Western blotting, with the permeability of *in vitro* GVB tested by Evans Blue-albumin assay. As shown in Figure 5A and 5C, compared with LPS control group, experimentally mild hypoxia (500 μM CoCl_2_ for 6 h) notably increased (*P* < 0.05) the expression of Claudin-1 protein in LPS-induced RIMVECs, and significantly decreased (*P* < 0.05) the content of EB-albumin in the lower chambers, suggesting that experimentally mild hypoxia (500 μM CoCl_2_ for 6 h) could repair the *in vitro* GVB disruption induced by LPS; whereas severe hypoxia (500 μM CoCl_2_ for 24 h) remarkably inhibited (*P* < 0.05) the expression of Claudin-1 protein in LPS-induced RIMVECs, and significantly increased (*P* < 0.05) the content of EB-albumin in the lower chambers, suggesting that severe hypoxia (500 μM CoCl_2_ for 24 h) could increase the *in vitro* GVB disruption induced by LPS. Meanwhile, as shown in Figure 5B and 5D, compared with TNF-α control group, experimentally mild hypoxia (500 μM CoCl_2_ for 6 h) significantly increased (*P* < 0.05) the expression of Claudin-1 protein in TNF-α-induced RIMVECs, and significantly decreased (*P* < 0.05) the content of EB-albumin in the lower chambers, suggesting that experimentally mild hypoxia (500 μM CoCl_2_ for 6 h) could also repair the *in vitro* GVB disruption induced by TNF-α; severe hypoxia (500 μM CoCl_2_ for 24 h) did not show any significant effect (*P* > 0.05) on neither Claudin-1 protein expression nor EB-albumin content in the lower chambers. These data suggest that experimentally mild hypoxia shows protective potential on *in vitro* GVB disruption under certain inflammatory microenvironment, while severe hypoxia may aggravate it.

**Figure 5.**
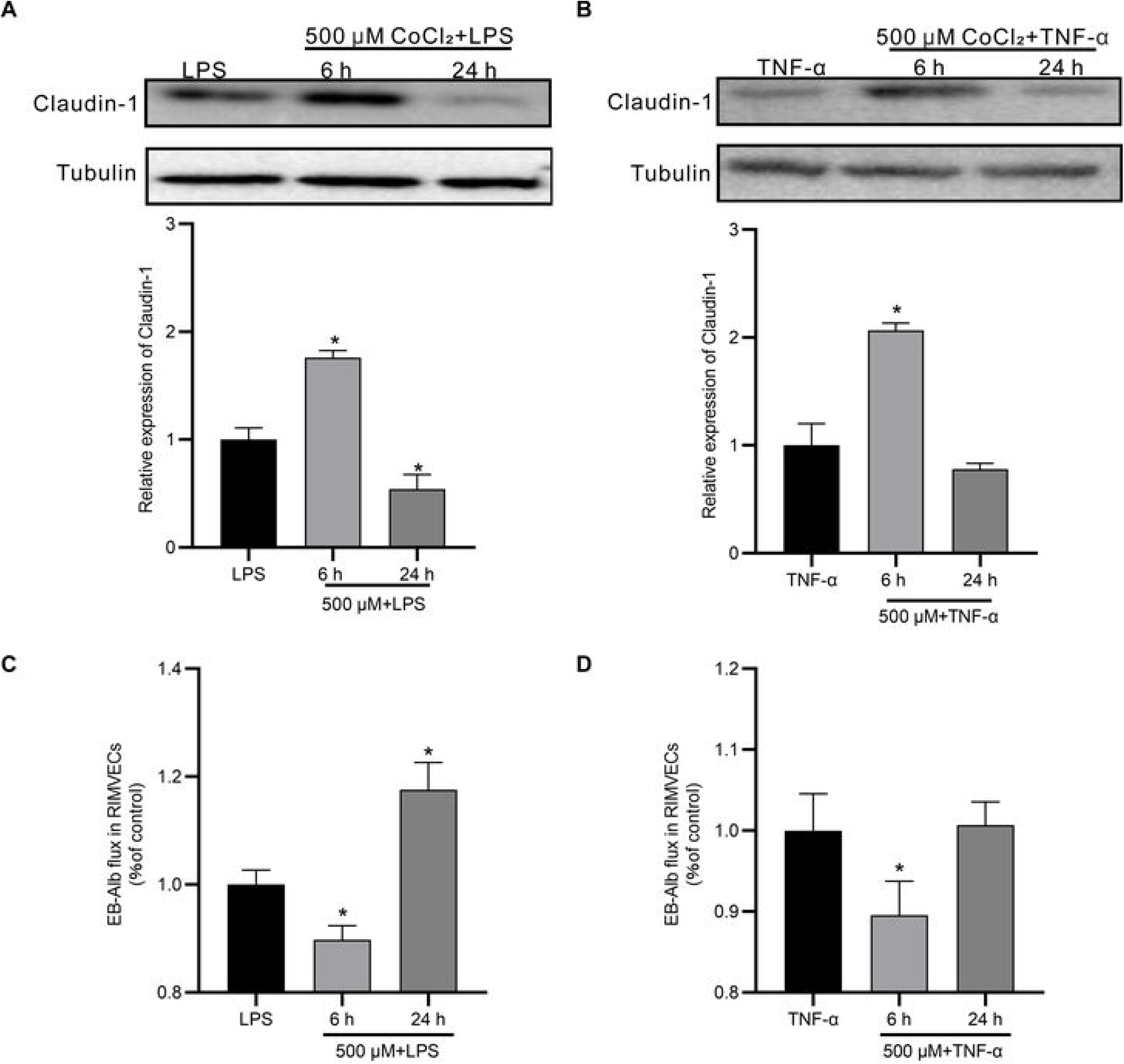
Hypoxia modulates inflammatory GVB disruption induced by LPS or TNF-α *in vitro*. (A) Western blotting analysis verified the effect of experimentally mild or severe hypoxia (500 μM CoCl_2_ for 6 h or 24 h) treatment on expression of Claudin-1 in RIMVEC-11 pre-treated by LPS (1 μM, 6 h) (n = 3). (B) Western blotting analysis verified the effect of experimentally mild or severe hypoxia (500 μM CoCl_2_ for 6 h or 24 h) treatment on expression of Claudin-1 in RIMVEC-11 pre-treated by TNF-α (0.3 μM, 6 h) (n = 3). (C) Effects of experimentally mild or severe hypoxia (500 μM CoCl_2_ for 6 h or 24 h) on the permeability of *in vitro* GVB (composed of RIMVEC-11) pre-treated by LPS (1 μM, 6 h). The permeability of the cell monolayer barriers was determined by EB-albumin efflux assay (n = 3). (D) Effects of experimentally mild or severe hypoxia (500 μM CoCl_2_ for 6 h or 24 h) on the permeability of *in vitro* GVB (composed of RIMVEC-11) pre-treated by TNF-α (0.3 μM, 6 h). The permeability of the cell monolayer barriers was determined by EB-albumin efflux assay (n = 3). Data information: In (A-D), data are presented as mean ± SD. \**P* < 0.05 was compared with the LPS or TNF-α group.

## Discussion

GVB is the deepest layer of protection in the intestine at the level of the gut vessel endothelium, representing the last obstacle to be crossed by a microorganism or a harmful toxin to reach the systemic circulation and organs far from the intestine^3^. Maintenance of GVB under various of intestinal microenvironment is essential to control the translocation of antigens into the bloodstream^2^. Intestinal hypoxia has an important role especially during acute intestinal inflammation in humans and mice, acting on several cells, including IECs and immune cells^18^. What are the consequences of hypoxia on GVB? Here, our results demonstrated the distinguishing bi-directionally regulatory mechanism of GVB induced by hypoxia in a HIF-1α-dependent manner *in vitro*.

The endothelial layer is the core structure of GVB^2,3^. It is located underneath the epithelial layer and is critical for the maintenance of water and protein balance between the intravascular and extravascular compartments in the intestinal system^20^. In the present study, CoCl_2_ was used as a chemically hypoxia inducer as previously described^21^, on the rat intestinal microvascular endothelial cells RIMVEC-11 and the GVB *in vitro*. The cell viability assay results showed that the RIMVECs were distinguishingly more susceptible to experimental hypoxia than the IECs were. Meanwhile, this tendency was also confirmed by the quantitative PCR results. Hypoxia-induced HIF-1α variation tendency at mRNA levels in IECs was much similar as that in RIMVECs, whereas evidently the degree of alteration in RIMVECs was much greater than that in IECs. Thereby, these results suggest that GVB may be susceptible to hypoxia. The GVB has morphological and functional characters similar to the BBB^2^. However, the GVB appears to be more permissive than the BBB because it allows the diffusion of larger molecules (up to 4 kDa)^3^. In the present study, the *in vitro* GVB model composed of RIMVECs was conducted. EB-albumin efflux assay results showed that experimentally mild hypoxia (500 μM CoCl_2_ for 6 h) could markedly strengthen the GVB function, whereas experimentally severe hypoxia (500 μM CoCl_2_ for 24 h) can lead the hyperpermeability and barrier disruption of GVB. In contrast, the IEB models, different from the GVB models, showed evident resistance under the same experimentally hypoxic conditions. Previous study has demonstrated that the intestinal epithelial barrier is uniquely resistant to changes elicited by hypoxia^22^. As a result of the unique anatomy, the intestinal epithelium shows a steep oxygen gradient at the steady state: the richly vascularized subepithelium reflects a high oxygenation level that decreases gradually to the almost anoxic lumen^15^. These results suggest that the GVB has a specifical characteristic of bi-directional regulation under hypoxia condition functionally. The intestinal mucosa adapts to fluctuations in blood flow toward the intestine (from 5% of total blood volume to 30% after a meal) that results in changes in local oxygen pressure^12^. In this way, it may be a regulatory adaption mechanism of the GVB under physiologically and pathologically hypoxic conditions. This characteristic of two-way regulation probably supports the hypoxia adaption of GVB under different stages of hypoxic conditions, to ensure the passage of luminal content and dietary antigens for tolerance induction while blocking bacterial translocation to the bloodstream.

The transcription factor HIF-1 functions as a key regulator of tissue barrier function and oxygen homeostasis^15,17,19,23,24^. To determine whether hypoxia regulates the GVB through HIF-1α and its mediated interendothelial TJs, we analyzed the protein expression and cellular distributions of HIF-1α, as well as the main TJ proteins including ZO-1, Occludin and Claudin-1 using Western blotting and immunofluorescence assay. Results suggested that hypoxia markedly activates HIF-1α to enter nucleus to play a regulatory role, and modulated intercellular TJ proteins in RIMVECs *in vitro*. Notably, we found that hypoxia-induced alteration of Claudin-1 protein was consistent with the trend of GVB permeability, which shares a similar regulatory pattern with that identified in a previous study that a epithelial HIF-1α / claudin-1 axis regulates barrier dysfunction in eosinophilic esophagitis^25^. Moreover, recovery of endothelial integrity after vascular injury is vital for endothelial barrier function and fluid homeostasis^26^,which is mostly driven by endothelial spreading and migration^27^. Hence, we assessed the cell cytoskeleton and endothelial migration of RIMVECs under hypoxia. The results suggested that the mild hypoxia could maintain and strengthen the GVB structurally at the endothelial level and activate the repair potentials of GVB injury, whereas the severe hypoxia disrupts the GVB structurally and inhibited the GVB repair potentials eventually. Collectively, these data confirm the distinguishing effects of hypoxia on GVB, as evidenced by the regulation of HIF-1α activation, interendothelial TJs, endothelial cytoskeleton, and endothelial migration and barrier repair. Therefore, to further confirm and elucidate the role of HIF-1α in hypoxia-induced modification of GVB, we downregulated HIF-1α expression in RIMVEC-11 by shRNA. The Western blotting results showed that the HIF-1α knockdown made Claudin-1 protein “blunted” to a certain extent in RIMVECs; the scratch assay results showed that HIF-1α mediated the regulation of hypoxia on GVB injury repair; the Evans Blue-albumin assay results showed that hypoxia regulated the permeability of GVB through a HIF-1α-dependent mechanism; the immunofluorescence assay results showed that the structurally destructive effects of severe hypoxia (500 μM CoCl_2_ for 24 h) on GVB, including abnormal reorganization and redistribution of cytoskeleton, wrinkled cell morphology and enlarged intercellular gaps, were all eliminated in sh-HIF RIMVECs. Therefore, these data above verify the key role of HIF-1α in hypoxia-induced modification of GVB. Previous studies have demonstrated the protective roles for the HIF-1 in both acute and chronic inflammatory diseases, particularly as they relate to mucosal surfaces involving epithelial cells^12,15,19, 28–30^. Also, HIF-1-dependent repression of adenosine kinase attenuates hypoxia-induced vascular leak^31^. However, some studies have shown that chronic HIF-1 stabilization can be harmful and exacerbate certain inflammatory conditions^18^. Here, we identified that hypoxia regulates the GVB functionally and structurally through a HIF-1α-dependent mechanism, but the actual effect on GVB (to strengthen or to destroy) may depend on the degree of hypoxia or HIF-1α activation. Physiologically, there is a gradient of O_2_ along the intestinal tissue as its concentration decreases more than ten times from the submucosa to the lumen^18^. Pathologically, the transmigrating neutrophils shape the mucosal microenvironment through localized oxygen depletion to influence resolution of inflammation^16^. In this way, the GVB in different locations may be affected by hypoxia during both homeostasis and active inflammation in different manners, demanding distinct adaptation.

The recovery of endothelial integrity after vascular injury is vital for endothelial barrier function and fluid homeostasis. Here, to further assess the effect of hypoxia treatment on GVB disruption, the inflammatory GVB disruption models were induced by LPS or TNF-α respectively. Both the Gram-negative bacterial LPS and the inflammatory cytokine TNF-α play crucial roles in the pathogenesis of the endothelial barrier disruption^32–34^, which are used as typical and representative inducements to conduct the barrier dysfunction model *in vitro*^20,32, 35–37^. In the present study, Western blotting and Evans Blue-albumin assay results showed that experimentally mild hypoxia showed protective potential on *in vitro* GVB disruption under certain inflammatory microenvironment, while severe hypoxia may aggravate it. Accordingly, certain hypoxia and HIF-1 activation might be considered as the putative therapeutic strategy to treat certain inflammatory and/or infectious conditions associated with GVB disorders.

In summary, our findings demonstrated that hypoxia could bi-directionally regulate the function of GVB through the HIF-1α-dependent mechanism *in vitro* for the first time. Furthermore, experimentally severe hypoxia could evidently exacerbate inflammatory GVB disruption induced by LPS or TNF-α, while the mild hypoxia may promote the repair. Our findings provide emerging evidence for the etiology and management of GVB disorders.

## Material and Methods

### Cells Cultures

The rat intestinal microvascular endothelial cell line RIMVEC-11 was established in our previous study^38^. RIMVEC-11 cells were cultured in Dulbecco’s modified Eagle’s medium (Thermo Fisher Scientific, Marina, USA) supplemented with 10% (vol/vol) fetal bovine serum (Gibco, Grand Island, USA), 100 U/mL penicillin G, and 100 μg/mL streptomycin at 37℃ in a humidified atmosphere of 5% CO_2_. The intestinal epithelial cell line IEC-6 was obtained from the National Infrastructure of Cell Line Resource. IEC-6 cells were cultured in Dulbecco’s modified Eagle’s medium (Thermo Fisher Scientific, Marina, USA) supplemented with 10% (vol/vol) fetal bovine serum (Gibco, Grand Island, USA), 100 U/mL penicillin G, 100 μg/mL streptomycin and 1 U/mL insulin (Y0001717, Sigma-Aldrich, USA) at 37℃ in a humidified atmosphere of 5% CO_2_. Chemically hypoxic conditions were incubated in a medium treated with 100 μM CoCl_2_ 6 h or 24 h, 500 μM CoCl_2_ 6 h or 24 h in a humidified atmosphere at 37°C.

### Cell Viability Assay

Cell viability was determined using a cell counting kit-8 (C0038, Beyotime, China). Briefly, RIMVEC-11 and IEC-6 were seeded at a density of 2 × 10^4^ cells per well in 96-well culture plates, respectively. After 18 h of cultivation, the cells were incubated with 100 μL of CoCl_2_ of different concentrations (0, 100, 200, 300, 400, 500, and 600 μM) for 6, 12, 24, and 48 h, respectively. After that, the cell viability was determined according to the standard protocols. The absorbance was measured at 450 nm using a Microplate Reader (BioTek Synergy H1, Agilent, USA).

### Establishment and Integrity Analysis of GVB and IEB *in vitro*

To establish an *in vitro* GVB model, RIMVEC-11 were seeded on transwell filters (Corning Inc, USA) at a density of 1.6 × 10^5^ cells per filter and grown as monolayers for 13 days grew as monolayers under conditions identical to that of RIMVEC-11. Similarly, to establish an *in vitro* IEB model, the IEC-6 cells were seeded on transwell filters at the same density and grew as monolayers under conditions identical to RIMVECs. Then, the endothelial cell monolayer barriers were incubated in the control mediums, 100 μM CoCl_2_ 6 h or 24 h, 500 μM CoCl_2_ 6 h (mild hypoxia), 500 μM CoCl_2_ 24 h (severe hypoxia). Then, the integrity of the cell monolayer barriers of RIMVEC-11 was determined by EB-albumin efflux assay as previously described^20,39^. Briefly, a 100 μL of buffer containing 4% bovine serum albumin (BSA, MW: 66.4 kDa) mixed with 0.67 mg/mL EB was added in the upper chambers. After 12 h of incubation, the cell culture medium (100 μL) was collected from the lower chambers. The Microplate Reader (BioTek Synergy H1, USA) measured the concentrations of EB-albumin in the collected samples. The EB flux of each group was expressed as a percentage relative to the control group.

### Preparation of Extracts of Nuclear and Cytoplasmic Proteins

The nuclear and cytoplasmic proteins were prepared strictly according to the Nuclear and Cytoplasmic Protein Extraction Kit (Beyotime, China) with about 1 × 10^7^ cells. The protein concentration was determined with a BCA protein assay (Beyotime, China).

### Western Blotting Assay

The cell samples were washed 3 times with PBS and lysed with RIPA buffer (Beyotime, China) on ice for 15 min. Protein concentrations were quantified by BCA protein assay (Beyotime, China). After that, the samples were separated by sodium dodecyl sulfate-polyacrylamide gel electrophoresis (SDS-PAGE) and transferred to polyvinylidene fluoride membranes (PVDF, Millipore, USA). The membranes were blocked with 5% low-fat milk for 2 h at room temperature (RT). After blocking with albumin from bovine serum for 1 h. They were incubated first with primary antibodies at room temperature and with specific primary antibodies (1:500-1000) at 4°C overnight. The membranes were reacted with primary antibodies, including anti-HIF1α (#36169), anti-Tubulin (#5666) (all from Cell Signaling Technology, USA), anti-ZO-1 (AF8394), anti-Occludin (AF7644), (all from Beyotime, China), anti-Claudin-1 (ab180158), anti-HDAC (ab280198) (all from Abcam, USA) The membranes were subsequently incubated with HRP-conjugated secondary antibodies (1:10000) for 40 min at room temperature. Blots were washed and incubated for 1 h with goat anti-rabbit or anti-mouse horseradish peroxidase-labeled antibodies (KPL, USA). The blots were revealed using ECL Plus Western Blotting Substrate (Thermo Fisher Scientific, USA). Images were captured by the Gel 3100 Chemiluminescent and Fluorescent Imaging System (Sage creation, China). The band density was quantified using Image J Software (version 1.51j8, National Institutes of Health (NIH), Bethesda, USA) with normalization to the Tubulin signal.

### Real-Time Quantitative PCR

Total RNA was extracted from the cell using TRIzol reagent (Thermo Scientific, USA), and first-strand complementary DNA (cDNA) was generated using a Reverse Transcription System Kit (TOYOBO, Japan) according to the manufacturer’s instructions. In this regard, each RT-qPCR reaction was conducted using three replicates with a final reaction volume of 20 μL per well, including 4 μL cDNA, 10 μL 2×SYBR Green Master mix (Vazyme, China), and 0.4 μL of upstream and downstream primers using an Mx3000P Real-Time PCR System (Agilent, USA). Primers used for PCR were as follows: *HIF-1α* F 5′-GCAACTGCCACCACTGATGA-3′, R 5′-GCTGCTTGAAAAAGGGAGCC-3′, *GAPDH* F 5′-GGATGCAGGGATGATGTTC-3′, R 5′-TGCACCACCAACTGCTTAG-3′, Each sample was run in triplicate, and a minimum of two independent experiments were performed for each sample.

### Immunofluorescence Assay

RIMVEC-11 were seeded onto sterile glass sliders in a 12-well culture plate at a density of 1.6 × 10^5^ cells per well. When the cell confluence on the glass sliders reached 90%, the cells were incubated in control mediums, 100 μM CoCl_2_ 6 h or 24 h, 500 μM CoCl_2_ 6 h (mild hypoxia), and 500 μM CoCl_2_ 24 h (severe hypoxia), the cells were mildly washed, fixed by 4% paraformaldehyde solution for 10 minutes, rinsed, and blocked by 5% BSA medium for 30 minutes at room temperature. Then, the cells were incubated with phalloidin (ab176756, Abcam) (1: 200) for 45 min, or the cell was added with the primary antibody HIF-1α (1:100) for 60 min at 37℃ and after triple wash in PBS, anti-mouse IgG, HRP-linked (#7076, Cell Signaling Technology) (1:2000) were subsequently added and incubated for 40 min at room temperature. Subsequently, the cells were washed 3 times before 4’,6-diamidino-2-phenylindole (DAPI) was added and rewashed. Representative images were captured by an Olympus FV3000 Confocal Laser Scanning Microscope (Tokyo, Japan).

### Knockdown of HIF-1α in RIMVEC-11 cells

To stably knockdown HIF-1α using jetPrime, a lentivirus particle was generated in HEK293T that was transfected with psPAX2 (packaging vector), pMD2.G (envelope vector), and pLKO.1 vector inserted with the sequence of shNC or shHIF-1α (shNC sequence 5′-TTCTCCGAACGT GTCACGT −3′, shRNAs targeting *HIF-1α* 5′-GCTACAAGAAACCGCCTATGA-3). A medium containing a lentivirus particle was harvested after three days of incubation and was centrifuged at 1,000 rpm for 5 min. The supernatants were filtered with 0.45-μm filter cells were transduced with shHIF-1α lentivirus in the presence of 5 μg/mL polybrene (Sigma-Aldrich, USA) for 72 h. The infected cells were transferred to a 100-mm culture dish and were selected for two weeks in a medium containing 3 μg/mL puromycin. The selected cell was named sh-NC or sh-HIF.

### Scratch Assay

RIMVEC-11 were seeded in 6-well culture plates to more than 95% confluency. Wounds were produced with a 200-μL pipet tip. After gently washing 3 times with PBS, cells were cultivated in DMEM containing 2% FBS with or without 100 μM CoCl_2_ and 500 μM CoCl_2_. Images of scratch gauge points were acquired at 0 h, 6 h, 24 h, and 48 h. post-scratching using a BX53 inverted microscope (Olympus, Japan). The size measure of the denuded areas was analyzed using ImageJ. The recovery rate (%) = (1 - the size measure of the uncovered space at post-scratching 0, 6, 12, 24 h, and 48 h time point/the size measure of the denuded area at the time point of 0 h) × 100.

### Statistical Analysis

Statistical analysis was performed using GraphPad Prism version 8.00 (GraphPad, CA, USA). All quantitative data were expressed as mean ± standard deviation (SD) for at least three separate occasions and analyzed by one-way variance (ANOVA), followed by the least significant difference (LSD) test for multiple comparisons. *P* values of less than 0.05 were considered statistically significant.

### Data Availability

The datasets generated during and/or analysed during the current study are available from the corresponding author on reasonable request.

## Acknowledgments

This work was supported by the National Natural Science Foundation of China (No. 31972656, 31902249, and 32102711), Key Research and Development Program of Zhejiang Province (No. 2023C02022), and ZAFU Research and Development Fund grant (No. 2021FR002).

## Author Contributions

PL and DW: Conceptualization, Data curation, Formal analysis, Methodology, Project administration, Software, Visualization, Writing -original draft, Writing-review & editing. JD and YZ: Investigation, Project administration. XW and HS: Funding acquisition, Supervision, Data curation. All authors contributed to refinement of the study protocol and approved the final manuscript.

